# Iconic Sound–Shape Correspondences in Aphasia

**DOI:** 10.64898/2026.05.18.725976

**Authors:** Josh Dorsi, Chaleece Sandberg, Simon Lacey, Lynne Nygaard, K. Sathian

## Abstract

**Purpose:** To examine speech iconicity for shape in aphasia, we compared shape iconicity ratings from people with aphasia to those from neurologically intact individuals and evaluated how these ratings relate to phonological and semantic processing profiles in aphasia.

**Method:** Twelve people with aphasia and twelve age- and gender-matched neurologically intact participants rated how rounded or pointed 50 auditory pseudowords sounded, using a 5-point scale. Ratings from participants with aphasia were compared to predicted shape iconicity ratings derived from reference ratings from prior work and to ratings from neurologically intact participants. For participants with aphasia, correlations between individual ratings and predicted ratings were related to measures of phonological and semantic processing.

**Results:** Ratings from people with aphasia were significantly correlated with both the predicted ratings and the ratings from neurologically intact participants. Relative to ratings from neurologically intact people, ratings from people with aphasia had weaker correlations to predicted ratings. There were indications that greater language impairment was associated with greater disruption of iconicity ratings; in particular, deficits in phonological segmentation and semantic processing were associated with reduced sensitivity to shape iconicity.

**Conclusion:** These findings suggest that sensitivity to shape iconicity is preserved to varying degrees in individuals with aphasia. Disruption of phonological and semantic processing appears to affect shape iconicity processing in aphasia.

---

Aphasia is a language disorder that affects approximately 2.5 million people in the US (Simmons-Mackie, 2018). Aphasia is most often caused by stroke, but can also be caused by traumatic brain injury or neurodegeneration. Estimates of the prevalence of post-stroke aphasia are variable, with an estimated median frequency of around 30% of stroke survivors (Flowers et al., 2016). People with aphasia (PWA) can struggle with the comprehension and production of oral and written language. Treatments for aphasia are not very efficient, typically involving many hours of one-on-one therapy for even modest improvement in language function (Fridriksson & Hillis, 2021). Innovative approaches to rehabilitation will be needed to meet the needs of the number of people living with aphasia, which is expected to rise as the population ages. Language iconicity, the general idea that the form of a word (in either spoken or sign language) intrinsically conveys its meaning, is an under-studied avenue that could support innovations in therapeutic approaches.

Speech iconicity is the association between the sound of a word and its meaning. For example, onomatopoeic words sound like the sounds they refer to (e.g., *bang*). Iconicity extends to non-auditory meanings as well; words such as *wiggle* and *crispy* also sound like their meanings (Winter et al., 2024). Iconicity is language-general, with similar sound-meaning relationships across different languages (Blasi et al., 2016), though not all of these relationships are universal and language-specific effects have been noted (Styles & Gawne, 2017). Additionally, pseudowords and real words have similar iconic sound-meaning correspondences (Sidhu et al., 2021; Kumar et al., 2025; Dorsi et al., 2026).

There is substantial evidence that iconicity plays a role in language learning. Some have proposed that sensitivity to sound-meaning correspondences forms the foundation for language acquisition (Imai & Kita, 2014). Iconicity-facilitated word learning can be observed in children as young as 14 months (Imai et al., 2015). Many of the earliest words learned by children are iconic (e.g., Sidhu et al., 2022), and adults tend to use iconic words when speaking to children (Perry et al., 2018). Infants prefer iconically congruent, relative to incongruent, speech-image pairs (Ozturk et al., 2013; Pena et al., 2011) and this congruency can facilitate word learning (e.g., Imai et al., 2015; see also Brand et al., 2018). The relationship between iconicity and word learning is even present in adults. Adults more accurately learned foreign language words that were paired with iconically congruent meanings than randomly assigned meanings (Nygaard et al., 2009; see also Revill et al., 2018 for an example with pseudowords).

A tantalizing prospect of the apparent link between iconicity and word learning is the suggestion that iconicity could be a rehabilitative tool for facilitating word-finding treatments in PWA. Yet, despite ample research investigating iconicity in the neurologically intact population, only two published reports have tested iconicity in PWA. Ammon et al. (1977) asked PWA to select which of a pair of abstract pictures corresponded to an auditory pseudoword. The pictures and pseudowords were selected from prior work that had established normative pseudoword- picture matches. Relative to these expected responses, PWA were less accurate than participants without aphasia, across speakers of three different languages. Moreover, accuracy correlated with language comprehension in the PWA. Although the performance of PWA was not explicitly compared to chance, it appears from this study that PWA have at least some residual capacity to assess iconicity.

More recently, Meteyard et al. (2015) asked PWA to perform a series of linguistic tasks using iconic (onomatopoeic) and non-iconic real words. PWA were significantly more accurate for iconic (compared to non-iconic) words in tasks that benefit from mapping phonology to semantic information. These researchers suggest that iconicity may support word processing in PWA through additional connections linking phonological forms, semantic representations, and modality-specific information, providing greater redundancy or alternative processing routes. Such an account suggests that iconicity may indeed be a strong tool for facilitating word retrieval in PWA.

To better understand iconicity in aphasia, we recently asked PWA and age- and gender- matched neurologically intact people (NIP) to classify auditory pseudowords as sounding either rounded or pointed, good or bad, big or small, and calming or exciting, in a series of two- alternative forced-choice tasks (Dorsi et al., 2024; Dorsi et al., 2025). The pseudowords in these tasks had previously been rated for each domain by young adult (aged 18-35), neurologically intact participants. Based on these prior ratings, half of the pseudowords in each task were chosen to be strongly associated with each response option. Consistent with Ammon et al. (1977), PWA were significantly less accurate than their neurologically intact counterparts. However, accuracy varied across domains. Importantly, for the shape (rounded/pointed) and arousal (calming/exciting) domains, PWA performed significantly better than chance, indicating that despite some disruption, iconicity is still present in PWA, consistent with Meteyard et al. (2015).

While our prior work indicated that iconicity is at least partially preserved in PWA, the extent of that preservation is still unclear. Ratings can provide more nuance to assessments of iconicity than binary judgments. The present work further examines the iconic domain of shape in PWA. We asked PWA and matched NIP to rate how rounded/pointed each of 50 auditory pseudowords sounded. We compared these ratings between groups and explored how these ratings correspond to standard aphasia assessments to test if the preservation of iconicity corresponds to language ability. We tested the hypothesis that disruption of iconicity would correspond to disruption of phonological and semantic processing in PWA.

## Method

### Participants

The ratings data from 12 PWA (7 female, 5 male) were included in this study. The age of this group ranged from 43 to 70 years (*M* = 55.6, *SD* = 8.0 years). All spoke American English as their first language (2 were multilingual). All developed aphasia following strokes and were 5 to 21 years post-onset (*M* = 10.8, *SD* = 4.8 years). Participants with aphasia were recruited based on self-reported aphasia diagnosis; aphasia type and severity were subsequently determined using the Western Aphasia Battery (WAB; Kertesz, 2006), administered during the course of the present study. Participant demographics are reported in Table 1.

**Table 1:** Gender was self-identified. Aphasia type was determined based on WAB assessment. ID denotes participant identifier.

| **ID** | **Gender** | **Age** | **Years Post Stroke** | **Aphasia Type** | **Aphasia Quotient** |
|---|---|---|---|---|---|
| 1 | F | 56 | 5 | Conduction | 68.6 |
| 2 | M | 53 | 6 | Anomic | 85.5 |
| 3 | M | 43 | 10 | Conduction | 70.7 |
| 4 | F | 63 | 12 | Anomic | 89 |
| 5 | F | 70 | 12 | Anomic | 81.8 |
| 6 | F | 61 | 6 | Anomic | 97.1 |
| 7 | M | 52 | 13 | Anomic | 94.1 |
| 8 | M | 54 | 15 | Conduction | 80.1 |
| 9 | M | 64 | 21 | Anomic | 87.2 |
| 10 | F | 57 | 15 | Anomic | 82.1 |
| 11 | F | 49 | 9 | Anomic | 92.6 |
| 12 | F | 45 | 6 | Anomic | 93.8 |

Twelve NIP (7 female, 5 male) were recruited to be age- and gender-matched to the PWA group. A gender-matched NIP with an age difference of less than 10 years was recruited for each PWA. This group ranged in age from 38 to 72 years (*M* = 55.3, *SD* = 11.6 years). All spoke American English as their first language (2 were multilingual) and had no history of neurological injury.

### Materials

The primary materials for this study were a set of 50 auditory consonant-vowel- consonant-vowel pseudowords, spoken by a female speaker whose first language was American English. Two independent judges listened to each pseudoword and confirmed they were spoken with neutral intonation and produced the intended phonemic content (see Table 2). These pseudowords were drawn from a larger set of stimuli used in prior research (McCormick et al., 2015; Lacey et al., 2020; Lacey et al., 2026) that asked young adult NIP to rate how rounded/pointed each pseudoword sounded. Based on these ratings, we selected pseudowords to span the rounded/pointed space, with 10 pseudowords selected to correspond to each of 5 categories: very rounded, moderately rounded, neither rounded nor pointed, moderately pointed, and very pointed (Table 2).

**Table 2.** Pseudowords grouped by predicted roundedness/pointedness ratings. Phonetic transcriptions are provided using the International Phonetic Alphabet (IPA).

| Very<br>Rounded | Moderately<br>Rounded | Neither | Moderately<br>Pointed | Very<br>Pointed |
|---|---|---|---|---|
| m□mo | d□uvu | s□t□u | kotu | t□et□i |
| mumo | l□me | sefe | v□zi | t□pi |
| molu | b□be | lile | t□□se | kIpe |
| mono | suso | leli | t□ku | tepe |
| molo | nIme | k□ko | vid□i | peti |
| m□lu | mene | sife | d□Ize | pite |
| m□mu | pupo | sIfe | z□ze | sit□e |
| l□mu | zovu | koku | d□ed□i | k□te |
| mulo | godo | gibe | bedi | t□ti |
| nonu | f□fu | fIfi | pIpe | kete |

### Procedure

This study consisted of three sessions completed on separate days. Each session was run remotely through video call using Zoom. Session 1 was a pseudoword rating task and was completed by all participants. Sessions 2 and 3 were only completed by participants with aphasia and involved standard aphasia assessments.

### Session 1

The rounded/pointed ratings of the auditory pseudowords were collected using a paradigm coded on PsychoPy software (Peirce 2007) which was run on the experimenter’s computer. The experimenter shared their computer’s screen and sound through Zoom. The paradigm began with on-screen instructions which informed the participants they would hear nonsense words, stating, “You will hear a nonsense word. This nonsense word is not a real word. Pretend the nonsense word is a real word (one you haven’t heard). Does it sound like it describes something rounded or pointed?” The instructions went on to explain that the participant would see a scale on the screen that they could use to indicate how rounded or pointed the pseudoword sounded. These instructions were read aloud by the experimenter, who confirmed that the participant understood the task before proceeding with the experiment. Participants were instructed to verbalize their responses. The participant’s intended response was confirmed as needed by the experimenter and entered into the computer.

Each trial began with a fixation cross (500 ms), after which the screen presented a visual scale from 1 to 5, labeling 1 as “Very Rounded”, 5 as “Very Pointed” and 3 as “Neither”. The “Very Rounded” and “Very Pointed” options included rounded and pointed abstract shapes (Figure 1) to assist with comprehension of the scale. These shapes had been rated as being strongly rounded or strongly pointed in a prior study (McCormick et al., 2015). There was a 500 ms inter-trial interval. The experiment began with 5 practice trials, during which the experimenter evaluated if the participant was experiencing any difficulty hearing the stimuli, was experiencing any other difficulty, and was performing the task. After the practice trials the experiment progressed to the main task in which the participant rated each of the 50 pseudowords once. The entire paradigm took about 20 minutes.

**Figure 1:**
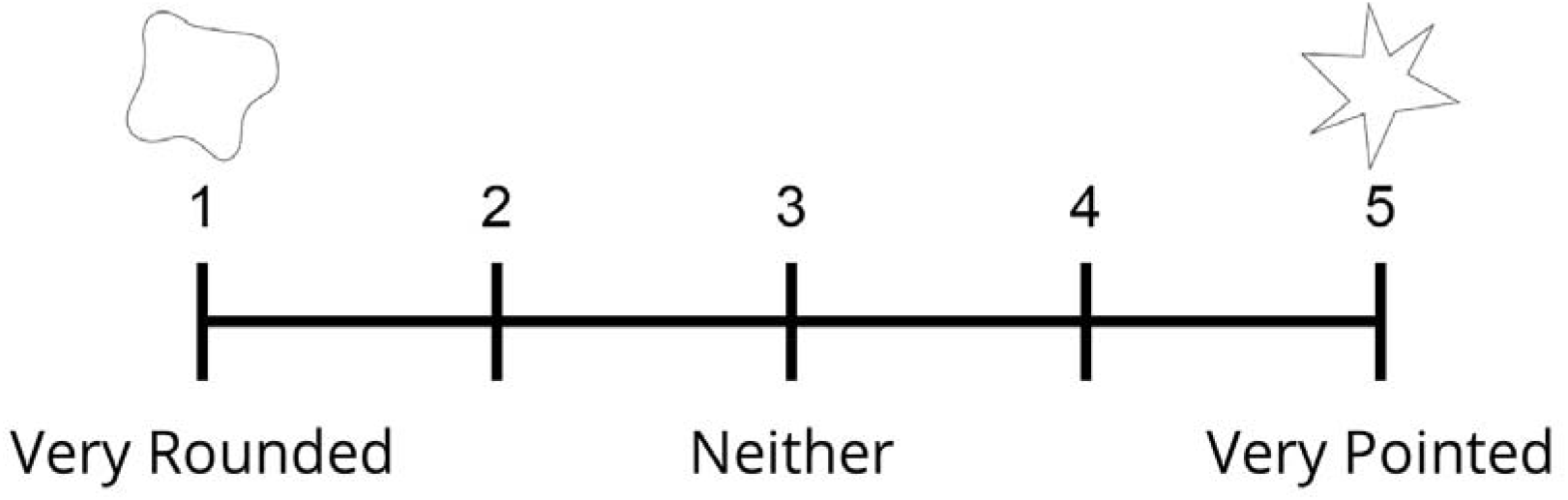
Response options for experiment. The scale was presented on participants’ screens while pseudowords were presented auditorily. Abstract shapes provide non-linguistic cues to the scale endpoints.

### Session 2

In session 2, which took about 3 hours, the WAB was administered. The WAB was used to classify the type of aphasia experienced by the participant and to calculate their aphasia quotient (AQ), which indicates aphasia severity.

### Session 3

Session 3 assessed PWA phonological processing with selected subtests of the Psycholinguistic Assessments of Language Processing in Aphasia (PALPA; Kay et al., 1992) and semantic processing with the 3-picture subtest of the Pyramids and Palm Trees Test (PAPT; Howard & Patterson, 1992). This session always followed the same sequence; participants completed phonological segmentation of initial sounds (PALPA subtest 16), then a same-different discrimination using nonword minimal pairs (PALPA subtest 1), then the three-picture variant of the PAPT, and ended with the phonological segmentation of final sounds (PALPA subtest 17). This schedule was chosen to minimize instruction confusion for the two phonological segmentation tasks.

Briefly, the phonological segmentation PALPA tasks asked the participants to select a letter presented onscreen that corresponded to the initial or final sound of a real word or pseudoword spoken by the researcher. The same-different discrimination PALPA task asked participants to indicate if a pair of pseudowords spoken by the researcher were the same or different. The PAPT task presented the participant with one picture centered above two other pictures; the participant was instructed to identify which of the two bottom pictures was more closely related to the one at the top.

### Analyses

We used representational similarity analysis (RSA; Kriegeskorte et al., 2008) to assess shape iconicity for our participants. Based on prior work, each pseudoword was coded from 1 to 5 on a rounded-to-pointed scale (Table 2). We generated a representational dissimilarity matrix (RDM) from the absolute difference between the predicted ratings for each pair of pseudowords (Figure 2). We next generated RDMs from the average ratings for the NIP and the average ratings for the PWA (Figure 2). The diagonal of an RDM—where each pseudoword is compared with itself—contains only zeros, and the matrix is symmetric about this diagonal. Spearman correlation (ρ) analyses were performed on the unique off-diagonal elements from one half of each RDM. In this way, we tested how similar the participant ratings for each group were to the predicted ratings, and how similar ratings were between the NIP and PWA groups.

**Figure 2:**
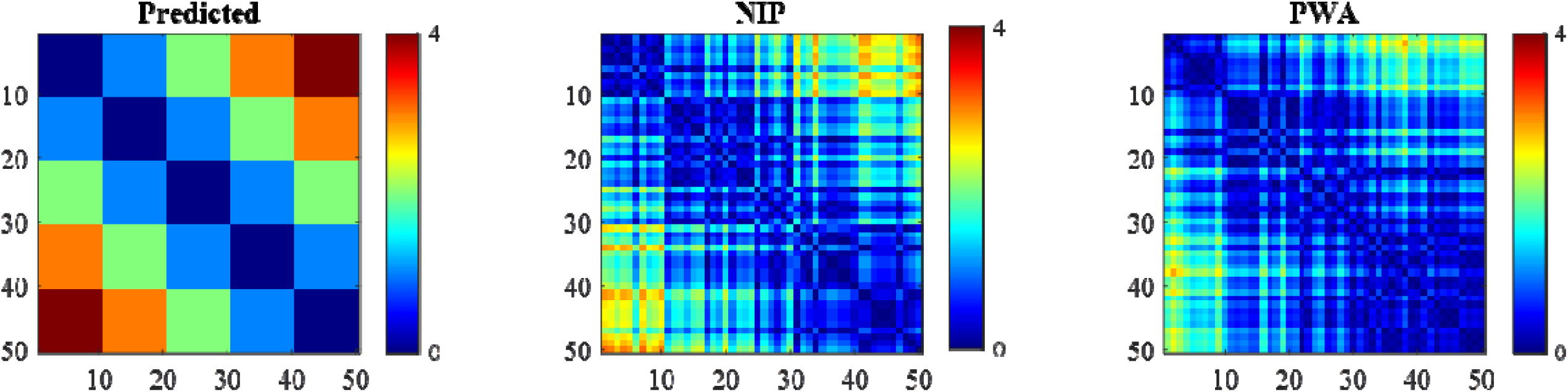
RDMs of predicted and group-averaged ratings. RDMs depict the pairwise differences in ratings across all pseudowords. Color indicates the absolute difference between ratings. The predicted RDM was derived from integer category values (1–5), whereas the NIP and PWA RDMs were derived from item ratings averaged across participants within each group.

In addition to the RDMs of group averages described above, we also generated an RDM for each participant following the same procedure. We conducted an RSA comparing each of these single-participant RDMs to the predicted RDM to analyze individual differences in iconicity. We converted the Spearman correlation coefficients to Fisher’s z (*Zr*) values and compared the NIP and PWA groups using Welch’s t-test.

Finally, we compared these individual RSAs (*Zr*) for the aphasia group to the assessments done in Sessions 2 and 3. Due to our limited sample size, we chose not to rely on inferential statistics for these comparisons. Instead, we report the relationships descriptively, focusing on the coefficient direction and magnitude rather than the corresponding *p*-value. To this end we adopt the benchmarks of .1, .3, and .5 for small, medium, and large effects, respectively (Cohen, 1992). Given the small sample size, any interpretation of these relationships is exploratory.

## Results

Unsurprisingly, the NIP group RDM was strongly correlated with the predicted RDM (ρ = .76, *p* < .001), demonstrating that ratings in the current work are consistent with those from prior work. Of more interest, the PWA group RDM was also found to be significantly correlated with the predicted RDM (ρ = .64, *p* < .001), indicating that speech iconicity is preserved in PWA. Moreover, the NIP and PWA RDMs were significantly correlated with each other (ρ = .46, *p* < .001), indicating that the sound-shape mappings used by PWA were similar to those of NIP.

While the *pattern* of ratings was similar for NIP and PWA groups, the groups differed significantly, as indicated by a Welch’s t-test (*t*[21.50] = 2.47, *p* = .02 [two-tailed], *d* = 1.01) of the *Zr* values. There were notable individual differences in how strongly each participant’s RDM correlated to the predicted RDM, with the NIP showing generally stronger correlations to the predicted RDM (Figure 3).

**Figure 3:**
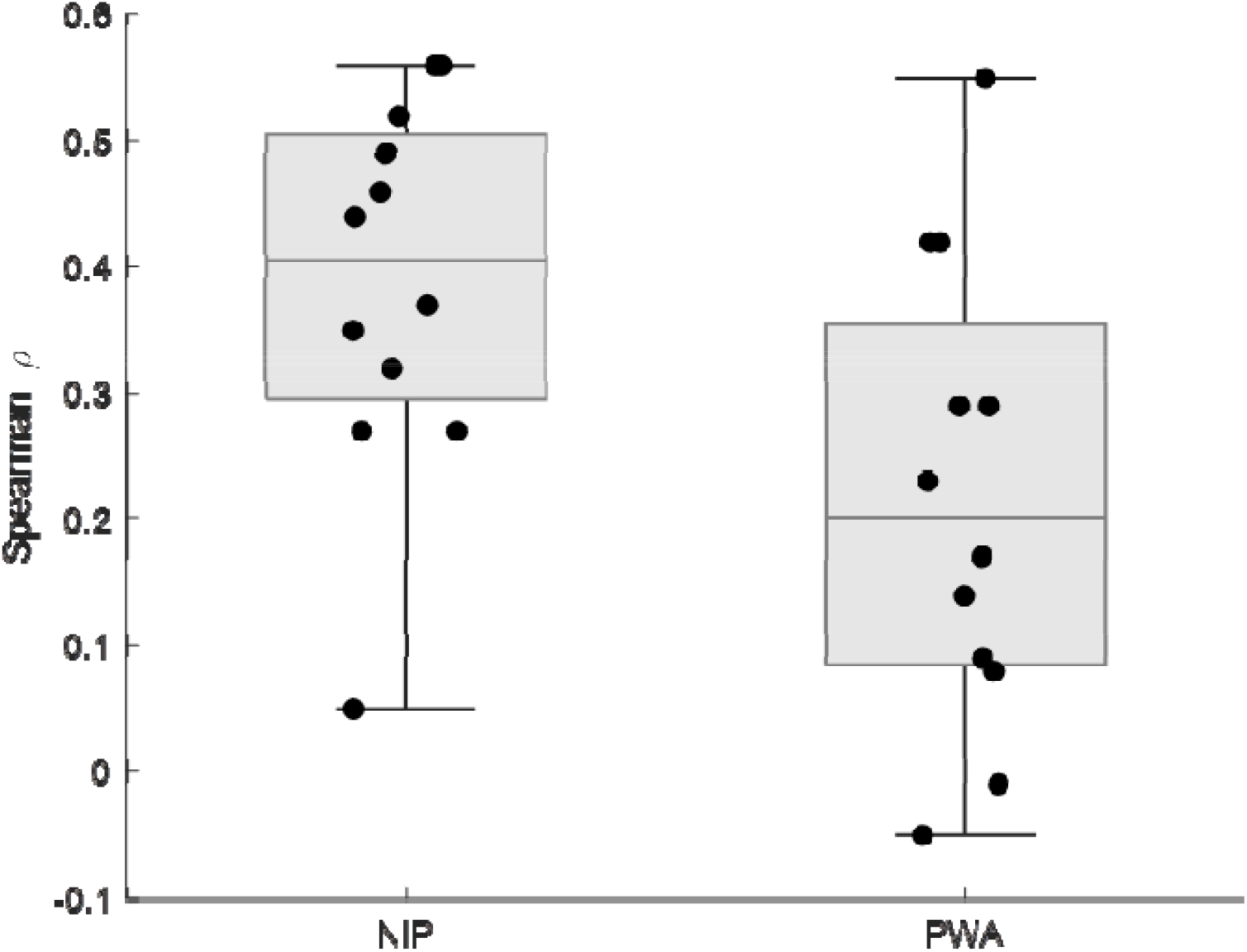
Individual Spearman correlations with the predicted RDM in NIP and PWA. Boxes show the interquartile range of Spearman correlations between each participant’s RDM and the predicted RDM for NIP and PWA; horizontal lines within the boxes indicate medians. Circles represent individual participants (jittered horizontally for visualization).

To examine the relationship between aphasia and iconicity, we compared each PWA’s *Zr* value (i.e., the Fisher transformed correlation between the individual’s RDM & the predicted RDM) to their aphasia assessment scores. The relationship between aphasia quotient (AQ) scores and *Zr* values (Figure 4) revealed a small positive trend (*r* = .15) indicating that less severe aphasia tended to be associated with stronger correlations to the predicted ratings.

**Figure 4:**
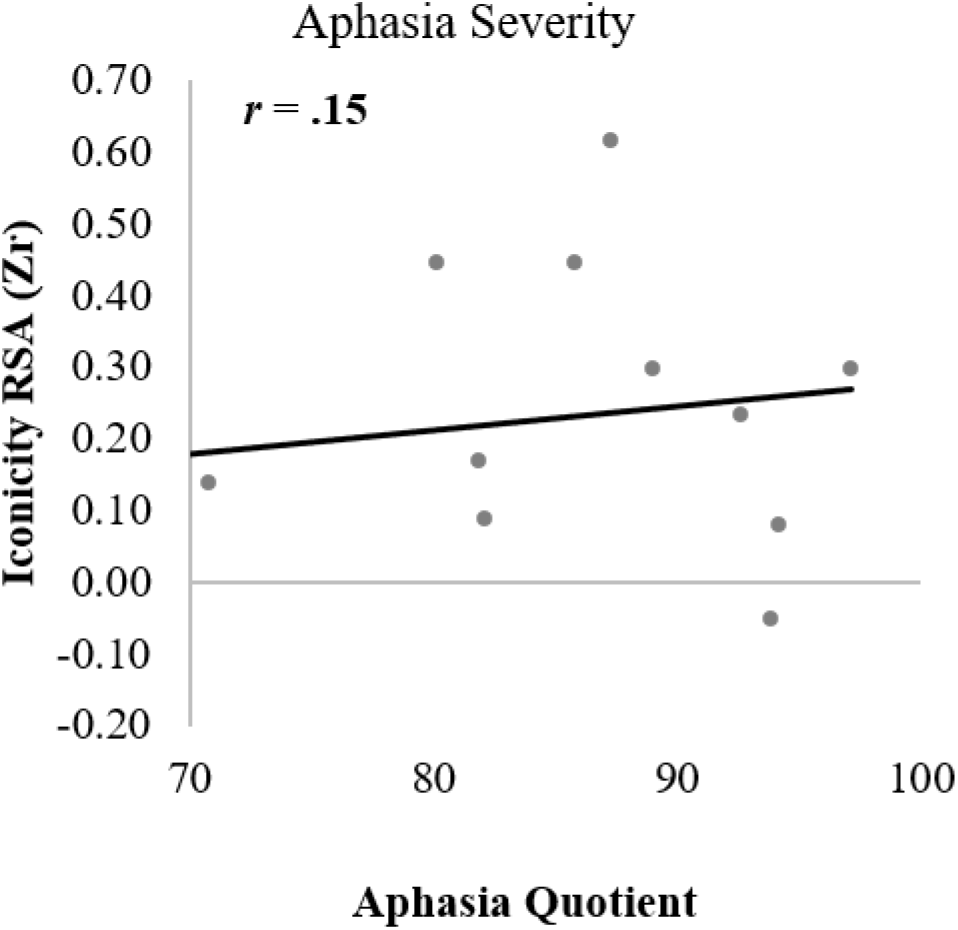
Iconicity-Related Zr Values by Aphasia Quotient. Shows *Zr* values of the Spearman correlations between each participant’s RDM and the predicted RDM plotted against aphasia quotient scores for each participant. Circles represent individual participants.

We next examined the relationship between phonological processing profiles and iconicity. Overall performance (*d*’) on same–different judgments of nonword pairs (PALPA 1) was not meaningfully related to *Zr* values (*r* = .06; Figure 5). This suggests that the ability to distinguish between different pseudowords was not associated with accuracy on the ratings task.

**Figure 5:**
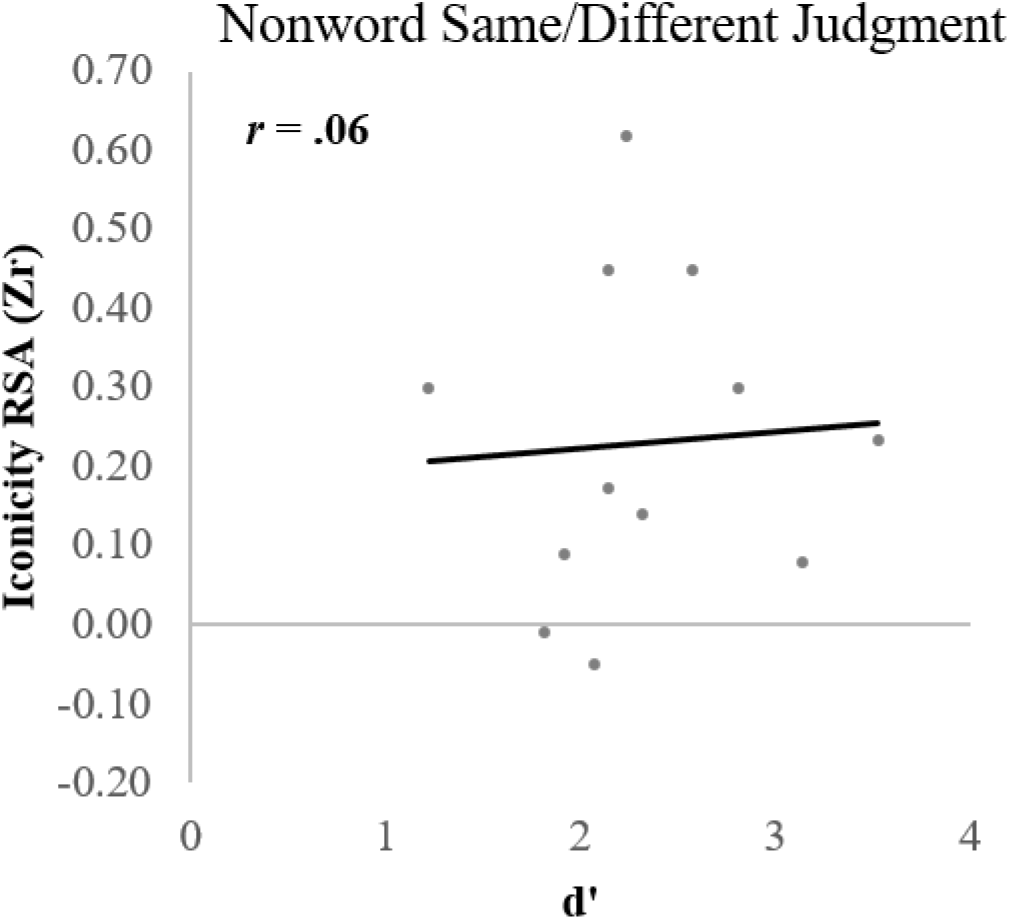
Iconicity-Related Zr Values and Phonological Discrimination Performance. Plots of iconicity-related *Zr* values against overall performance (d’) on the PALPA 1 same–different nonword discrimination task (averaged across same and different types).

To test whether phonological ability was related to iconicity ratings, we next examined the accuracy of phonological segmentation of initial and final segments of nonwords (PALPA 16 & 17, respectively). Individual *Zr* values were strongly associated with the ability to segment initial nonword sounds (PALPA 16; *r* = .65). In contrast, segmentation for final-position segments was only weakly related to individual *Zr* values (PALPA 17; *r* = .22). These trends are summarized in Figure 6.

**Figure 6.**
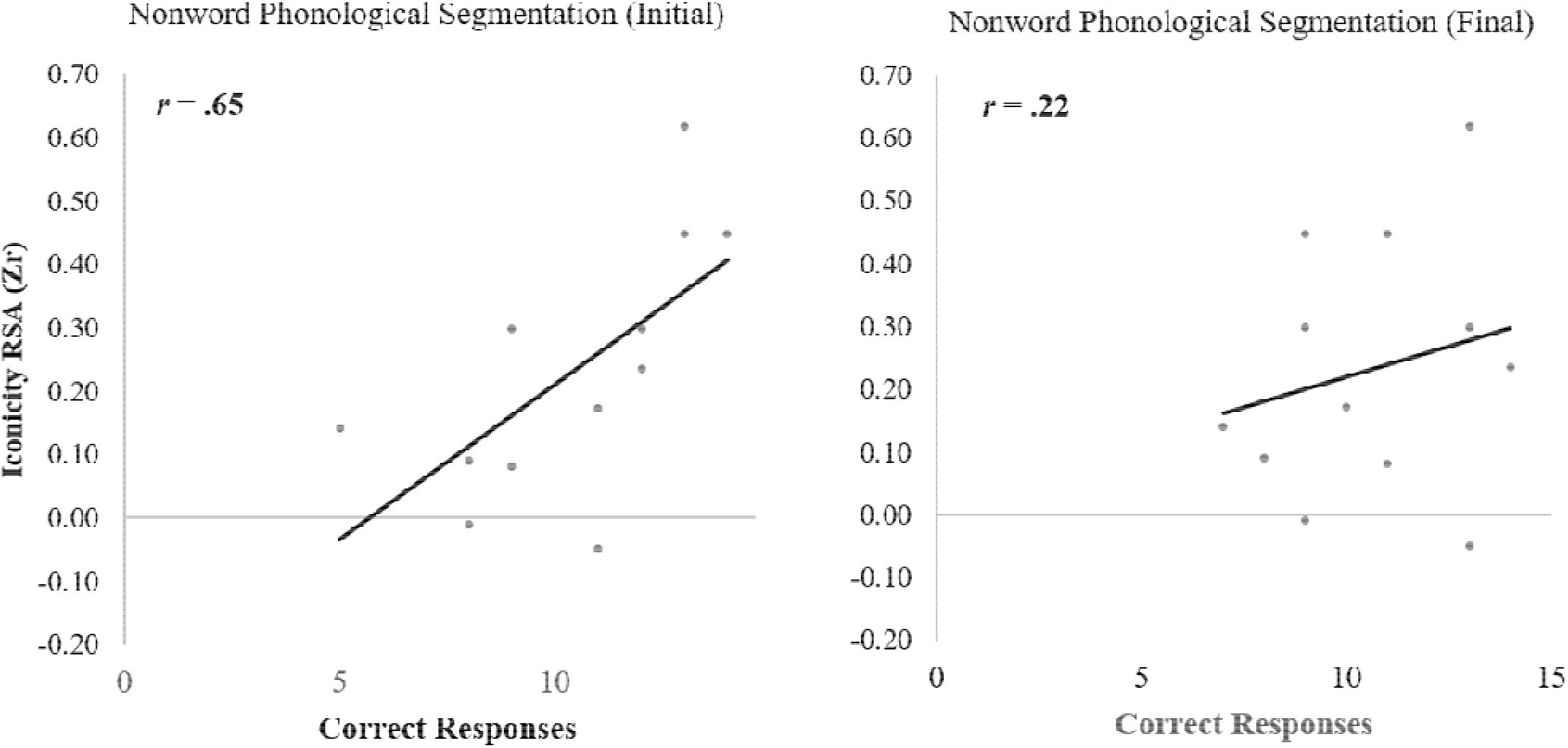
Iconicity-Related Zr Values and Position Phonological Segmentation. Iconicity-related *Zr* values against performance on phonological segmentation of initial segments (PALPA 16, left panel), and final segments (PALPA 17, right panel). Each circle represents an individual participant.

Finally, we found a strong correlation between individual *Zr* values and performance on the PAPT (*r* = .57; Figure 7), indicating that PWA with better semantic processing had stronger correlations between their ratings and the predicted ratings.

**Figure 7.**
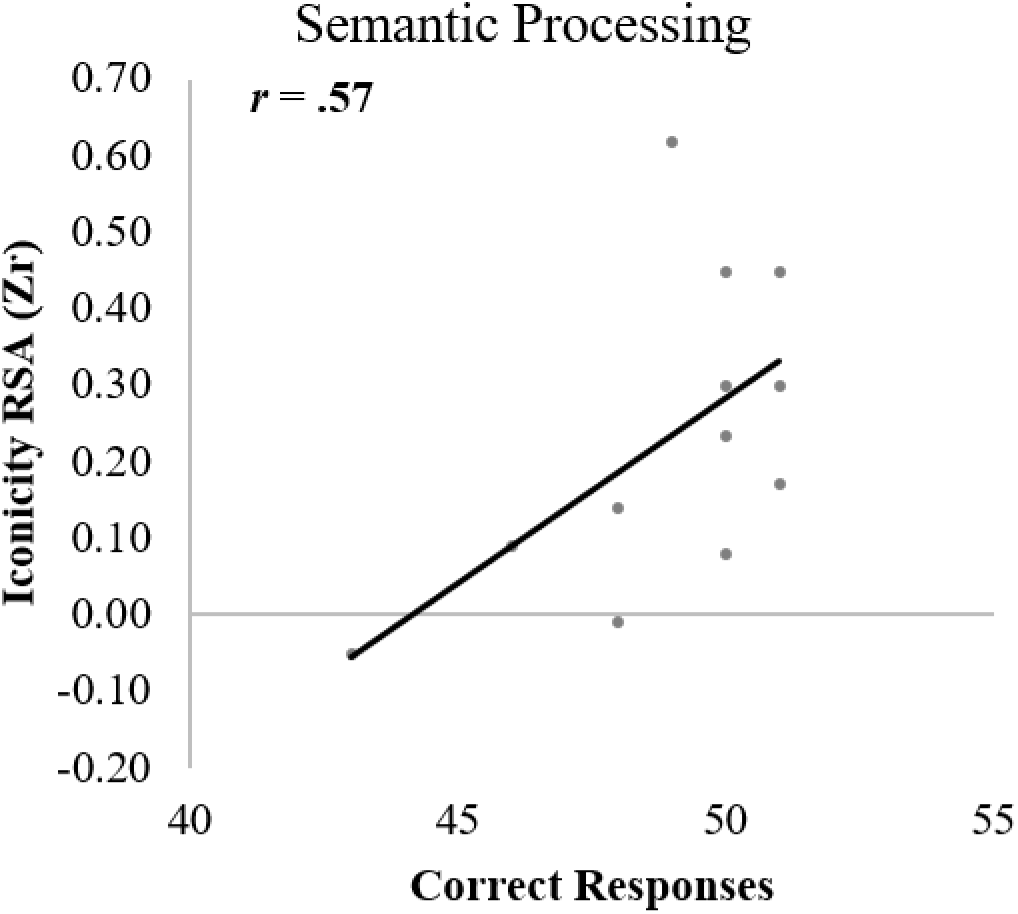
Iconicity-Related Zr Values and Semantic Processing. Iconicity-related *Zr* values against performance on the PAPT, a measure of semantic processing. Each circle represents an individual participant.

## Discussion

We investigated whether shape iconicity is preserved in PWA by comparing their rounded/pointed ratings of auditory pseudowords to ratings from NIP. We found that the rounded–pointed sound ratings produced by PWA were positively correlated with the ratings of NIP, providing evidence that iconic sound–meaning mappings are preserved in aphasia. A group-level comparison of individuals’ *Zr* values (i.e., how strongly their ratings correlated with ideal ratings) showed a statistically significant difference, indicating weaker iconicity in PWA relative to NIP, consistent with earlier findings reported by Ammon et al. (1977).

Importantly, the pattern of individual *Zr* values further indicates that the degree to which iconicity is preserved in aphasia may be related to the nature and severity of the language impairment. While individuals with more severe language disruption tended to show lower *Zr* values (i.e., weaker correspondences between their ratings and the predicted ratings), this relationship was weak. However, individual differences in *Zr* values were more strongly associated with differences in phonological and semantic processing abilities, suggesting that sensitivity to iconic sound–meaning structure may depend on specific linguistic resources that are variably compromised in aphasia.

Given the small sample size and the limited variety of aphasia types sampled, inferences concerning the relationship between aphasia type and performance on the pseudoword rating task are limited. It should also be acknowledged that aphasia type may not accurately capture language-system disruption experienced by PWA (Tremblay & Dick, 2016). Three PWA were classified as having conduction aphasia, with the remainder classified as having anomic aphasia. The conduction group trended toward lower *Zr* values, but two of the three participants also exhibited the lowest overall language ability as indexed by the AQ. Further work with larger and more diagnostically balanced samples will be required to clarify the relationship between aphasia type and iconicity.

Aphasia severity appeared to be more closely related than categorical diagnosis to *Zr* values. As shown in Figure 4, individuals with less severe aphasia tended to show stronger correspondence with the predicted ratings, though even this relationship is weak. This pattern aligns with findings reported by Ammon et al. (1977), who observed that accuracy in a pseudoword–shape matching task was correlated with overall language performance in PWA.

Several accounts propose that iconic sound–meaning correspondences provide an important foundation for early word learning (e.g., Imai & Kita, 2014). Consistent with this view, prior studies have shown that children’s earliest words tend to be relatively iconic (Sidhu et al., 2022), that learners acquire pseudoword–image mappings more rapidly when they follow iconic phonological patterns (Imai et al., 2015), and that adults learn and retain iconic meanings of foreign words more effectively than arbitrary mappings (Nygaard et al., 2009; Revill et al., 2018). If iconicity can scaffold word learning, it may also represent a potential resource for supporting lexical retrieval in aphasia. Given that PWA in the present study exhibited measurable, albeit somewhat reduced, sensitivity to iconic sound–meaning structure as indexed by their *Zr* values, future work could explore whether iconic cues might be incorporated into established word-finding therapies to support language rehabilitation.

## Understanding Iconicity

A persistent challenge in research on iconicity concerns the source of the phonological– meaning correspondences observed for both real words and pseudowords. Specifically, it remains unclear to what extent such mappings reflect regularities extracted from existing lexical items and subsequently generalized to pseudowords, or whether they arise from prelexical sound–meaning associations that influence both words and pseudowords (Sidhu, 2025). Although the present results do not adjudicate between these accounts directly, the relative strength of the associations between individual *Zr* values and the various phonological and semantic measures may offer some insight into the representations that support iconic sound– meaning correspondence.

The strongest associations with *Zr* values were observed for tasks assessing phonological segmentation ability. The relatively strong correlations between *Zr* values and segmentation performance suggest that access to internal phonological structure plays an important role in iconic sound–meaning correspondence in aphasia. Because performance on these tasks may be influenced by phonological–orthographic mapping abilities, we conducted *post hoc* partial correlations controlling for nonword spelling performance on the WAB (see Supplemental Analyses). Importantly, these partial correlations were even stronger, lending further support to the interpretation that phonological segmentation ability—and not orthographic knowledge per se—is closely related to sensitivity to iconic sound–meaning structure. This interpretation converges with recent findings from young adult NIP, which demonstrate that phoneme identity is a significant contributor to iconicity ratings of pseudowords (Lacey et al., 2026; Westbury et al., 2018; Monaghan & Fletcher, 2019).

In contrast to the segmentation tasks, the same–different judgment task had no relationship to *Zr* values. This discrepancy may reflect differences in the phonological demands of these tasks. Same–different judgments can be performed based on auditory or phonetic cues and do not necessarily require explicit phonemic identification. It is also possible that the phonological contrasts tested in some same–different conditions are not systematically aligned to shape iconicity.

In addition to phonological processing, we also observed a fairly large association between *Zr* values and performance on the PAPT, suggesting that iconic sound–meaning correspondence may also relate to semantic processing ability. Because the PAPT is a picture-matching task with minimal (if any) phonological demands, this association indicates that iconicity may engage semantic access that is disrupted in aphasia. However, it is important to note that most participants performed near ceiling on this measure, limiting variability and constraining the strength of inferences that can be drawn (see Supplemental Analysis for sensitivity analyses and related post hoc checks). Future work including a broader range of semantic abilities will be necessary to clarify the nature of this relationship.

Previous studies have reported advantages for iconic over non-iconic words in lexical decision tasks in both NIP (Sidhu et al., 2020) and PWA (Meteyard et al., 2015). These effects have been interpreted as reflecting enhanced phonological–semantic connectivity for iconic words within the lexical system, although it is also possible that iconicity may primarily facilitate task-specific decision processes rather than lexical access per se (Sidhu, 2025). The present findings—that individual *Zr* values (the correlation between a participant’s ratings and predicted ratings) are related to phonological segmentation and semantic processing—are more consistent with an account in which iconicity reflects structured phonological–semantic mappings within the lexical system. Together, these results suggest that iconic sound–meaning correspondence draws on linguistic representations that are variably compromised in aphasia.

## Limitations

This work has several limitations that also motivate directions for future research. First, the modest and relatively homogeneous sample of 12 participants necessitates cautious interpretation of all reported results. One consequence of samples of this size is that individual participants can exert a disproportionate influence on the observed associations. For example, the correlation between overall aphasia severity and *Zr* values is notably impacted by a single participant who had the third highest AQ and the lowest *Zr* value. Excluding this participant from the statistical analysis increased the correlation between AQ and *Zr* value, from weak to moderate. The impact of removing this participant from the other analyses is reported in more detail in the Supplemental Analyses. Given the present sample size, it is difficult to determine whether this participant represents an unusual case or an initial observation of a broader pattern. Until additional data are collected, both of these possibilities must be considered when evaluating the reported results.

Finally, the present experiment examined only iconic shape ratings of pseudowords, on a rounded–pointed dimension, and thus targeted a relatively narrow domain of iconicity. Although this dimension has been widely studied and offers a useful starting point, iconic sound–meaning correspondences extend across multiple semantic domains. Determining whether similar relationships between iconicity, phonological processing, and semantic processing are observed for other types of iconic mappings will be essential for developing a more general account of iconic sound–meaning relations in aphasia. Addressing these limitations will be critical both for refining theoretical models of iconicity and for evaluating its potential role in supporting language recovery.

## Supporting information

Supplemental Analyses

## Acknowledgments

This work was supported by institutional funds provided to KS by Penn State College of Medicine.

An AI-based language tool assisted in this work. Portions of the text were edited for clarity and style by Microsoft Copilot. The authors take full responsibility for the content. An AI- based language tool (Microsoft Copilot) produced portions of the code and command syntax for generating figures and handling data. All code was reviewed, validated, and executed by the first author, who confirmed the accuracy of each step and the integrity of the resulting data and figures.

## Data Availability Statement

The datasets generated during and/or analyzed during the current study are available in the Open Science Framework repository, https://osf.io/tevhc/overview?view_only=9b7f7fa9d43b4d99a72c29f90192579c.

