## Supplemental Analyses for "Iconic Sound–Shape Correspondences in Aphasia"

**Phonological Segmentation and Orthographic Mapping**

Performance on the phonological segmentation tasks of the PALPA (16 and 17) may be influenced by phonological–orthographic mapping ability. Given the limited sample size, it was not possible to fully control for phonological-to-orthographic mapping skills in the primary analyses. To provide additional context for the results reported in the main text, Table S1 presents individual participant data from the PALPA 16 and 17, alongside Supplemental Reading and Writing scores from the WAB and iconicity *Zr* values derived from the ratings RSA.

We additionally examined partial correlations between phonological segmentation performance and ratings-based iconicity correlations while controlling for nonword writing performance on the WAB. These relationships remained robust: partial correlations were strong for initial phoneme segmentation (PALPA 16; *r* = .68) and moderate for final phoneme segmentation (PALPA 17; *r* = .38).

**Sensitivity Analysis**

With our limited sample size, individual participants can have an outsized impact on the statistical analysis. To evaluate this impact, we excluded the PWA participant with the smallest *Zr* value (indicating the weakest alignment to ideal iconic ratings) and their matched control from each of the main analyses.

The correlation between the PWA group RDM and the predicted RDM remained robust (ρ = .65, *p* < .001) as did the correlation between the PWA group RDM and the NIP group RDM (ρ = .45, *p* < .001). The Welch’s t-test was also consistent with that reported in the main text (*t*[19.93] = 2.19, *p* = .04 [two-tailed], *d* = 0.93). Thus, we conclude that the participant with the lowest iconicity-related *Zr* value did not have a substantial impact on the overall iconicity ratings of the PWA group.

The changes to the correlations between the *Zr* values and aphasia assessments as a result of this sensitivity analysis are summarized in Table S2. Notably, the sensitivity analysis suggests that the relationship between overall language function, indexed by the AQ, and iconicity might actually be stronger than indicated in the analyses reported in the main text. Additionally, the largest change in the sensitivity analysis was the association between *Zr* values and PALPA 17 performance, which increased to a moderate correlation, consistent with our interpretation of a role for phonological segmentation in iconicity. In contrast, the sensitivity analysis indicates that the relationship between semantic processing, indexed by the PAPT, might be weaker than indicated by the analysis reported in the main text. As there are theoretical implications of a relationship between semantic processing and iconicity, this deserves careful consideration, which is provided in the next section.

**Post-hoc Evaluation of Semantic Processing**

In addition to having the lowest *Zr* value, the participant removed for this sensitivity analysis also had the lowest score on the PAPT. This is notable because this participant also had a relatively high AQ. The AQ includes measures of semantic processing, so a dissociation between it and the PAPT warrants caution. Indeed, the correlation between the AQ and PAPT performance was moderate in the sensitivity analysis (*r* = .45) but near zero in the analysis with the full sample (*r* = .04). It is possible that this participant’s performance on the PAPT reflected factors unrelated to semantic processing.

As an alternative test of the relationship between semantic processing and iconicity, we took advantage of the Word Fluency component of the WAB. In this task, the participants name as many animals as they can in one minute. The correlation between word fluency and *Zr* values was moderate in both the full sample (*r* = .35) and the sensitivity analysis (*r* = .34; Figure S1).

**Table S1.** Individual participant performance on phonological segmentation tasks from the PALPA (16: initial phoneme segmentation; 17: final phoneme segmentation) for both nonwords and real words, Supplemental Reading and Writing subtests of the Western Aphasia Battery (WAB), and the iconicity *Zr* values derived from the ratings representational similarity analysis (RSA). ID denotes participant identifier.

|  | **NonWords** | | | | **Words** | | | |  |
| --- | --- | --- | --- | --- | --- | --- | --- | --- | --- |
| **ID** | **Writing** | **Reading** | **Initial** | **Final** | **Writing** | **Reading** | **Initial** | **Final** | **Zr** |
| **1** | 0 | 0 | 8 | 9 | 3 | 7 | 23 | 14 | -0.01 |
| **2** | 2 | 6 | 13 | 11 | 3 | 10 | 27 | 23 | 0.45 |
| **3** | 0 | 1 | 5 | 7 | 0 | 7 | 11 | 14 | 0.14 |
| **4** | 1 | 5 | 9 | 9 | 6 | 10 | 26 | 20 | 0.30 |
| **5** | 0 | 5 | 11 | 10 | 0 | 10 | 27 | 18 | 0.17 |
| **6** | 5 | 6 | 12 | 13 | 10 | 10 | 26 | 30 | 0.30 |
| **7** | 6 | 7 | 9 | 11 | 7 | 10 | 22 | 27 | 0.08 |
| **8** | 0 | 7 | 14 | 9 | 7 | 9 | 26 | 21 | 0.45 |
| **9** | 0 | 9 | 13 | 13 | 4 | 9 | 26 | 24 | 0.62 |
| **10** | 1 | 2 | 8 | 8 | 0 | 10 | 20 | 24 | 0.09 |
| **11** | 4 | 8 | 12 | 14 | 7 | 10 | 27 | 26 | 0.23 |
| **12** | 2 | 10 | 11 | 13 | 7 | 10 | 27 | 26 | -0.05 |

**Table S2.** Changes between the main analysis of the full sample of PWA (N = 12) and the sensitivity analysis in which the person with the lowest accuracy on the iconicity ratings task (*Zr* value) was removed from the correlation tests (N = 11). Pearson correlations between the *Zr* values and each of five assessments of language processing for the main analysis and the sensitivity analysis are reported.

| **Assessment** | **Full sample** | **Sensitivity analysis** | **Δr** | **Interpretation** |
| --- | --- | --- | --- | --- |
| **AQ** | .15 | .33 | .18 | Increases: Weak to Moderate |
| **PALPA 1** | .06 | < .01 | ~.05 | No meaningful change |
| **PALPA 16** | .65 | .76 | .11 | No meaningful change |
| **PALPA 17** | .22 | .44 | .22 | Increases: Weak to Moderate |
| **PAPT** | .57 | .41 | .16 | Decreases: Strong to Moderate |

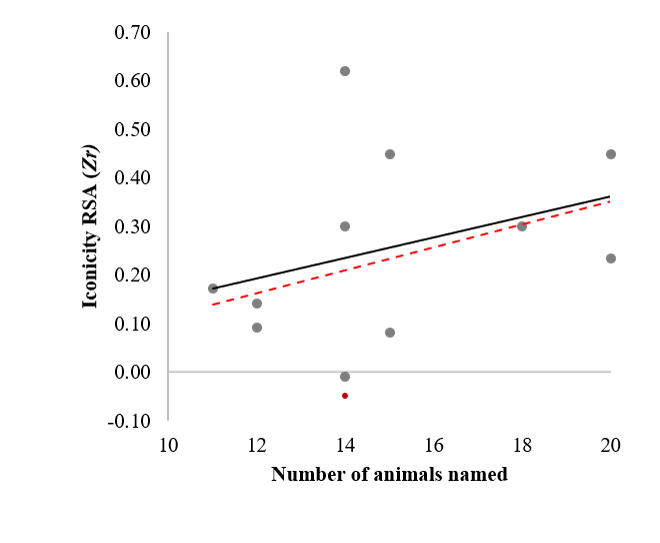

**Figure S1:** *Iconicity Ratings and Semantic Processing*. The vertical axis plots the iconicity-related *Zr* values, the horizontal axis plots the number of animals named in the word fluency task of the WAB. The dark points and trend line correspond to the N = 11 sensitivity analysis, the red point and line correspond to the N = 12 analysis.
